# Performance of Abbott ID NOW COVID-19 rapid nucleic acid amplification test in nasopharyngeal swabs transported in viral media and dry nasal swabs, in a New York City academic institution

**DOI:** 10.1101/2020.05.11.089896

**Authors:** Atreyee Basu, Tatyana Zinger, Kenneth Inglima, Kar-mun Woo, Onome Atie, Lauren Yurasits, Benjamin See, Maria E. Aguero-Rosenfeld

## Abstract

The recent emergence of the SARS-CoV-2 pandemic has posed formidable challenges for clinical laboratories seeking reliable laboratory diagnostic confirmation. The swift advance of the crisis in the United States has led to Emergency Use Authorization (EUA) facilitating the availability of molecular diagnostic assays without the more rigorous examination to which tests are normally subjected prior to FDA approval. Our laboratory currently uses two real time RT-PCR platforms, the Roche Cobas SARS-CoV2 and the Cepheid Xpert Xpress SARS-CoV-2. Both platforms demonstrate comparable performance; however, the run times for each assay are 3.5 hours and 45 minutes, respectively. In search for a platform with shorter turnaround time, we sought to evaluate the recently released Abbott ID NOW COVID-19 assay which is capable of producing positive results in as little as 5 minutes. We present here the results of comparisons between Abbott ID NOW COVID-19 and Cepheid Xpert Xpress SARS-CoV-2 using nasopharyngeal swabs transported in viral transport media and comparisons between Abbott ID NOW COVID-19 and Cepheid Xpert Xpress SARS-CoV-2 using nasopharyngeal swabs transported in viral transport media for Cepheid and dry nasal swabs for Abbott ID NOW. Regardless of method of collection and sample type, Abbott ID NOW COVID-19 had negative results in a third of the samples that tested positive by Cepheid Xpert Xpress when using nasopharyngeal swabs in viral transport media and 45% when using dry nasal swabs.

## Introduction

The novel coronavirus, severe acute respiratory syndrome coronavirus 2 (SARS-CoV-2) causing coronavirus disease-2019 (COVID-19) was first identified in Wuhan Jinyintan Hospital, China in December 2019. Since then, it has spread widely, reaching the status of global pandemic as declared by the World Health Organization on March 11^th^, 2020 (1). As of May 25th, 2020, over 5 million cases and over 342 thousand deaths have been reported worldwide (2) while in New York City, there were over 196 thousand reported cases and more than 16 thousand deaths (3, 4).

SARS-CoV-2 virus is detected in various human samples including respiratory and fecal, but testing has been validated for PCR primarily for nasal, nasopharyngeal, and oropharyngeal specimens. Positive samples for SARS-COV-2 were found in Chinese patients 1-2 days prior to symptom onset and persisted up to 2 weeks in severe cases (1).

While long-term mitigation and prevention goals include prognostic markers, therapeutics, and vaccines (1), the urgency of this rapidly developing crisis in the United States has prioritized the immediate goal of diagnostic testing, advanced through Emergency Use Authorization (EUA) by the FDA (5, 6). The initiative to expand and accelerate testing has eased the usual scrutiny that new assays would normally undergo prior to release. This obligates clinical laboratories to more carefully assess the overall performance of the various rapidly emerging testing platforms prior to implementation. Several Nucleic Acid Amplification tests (NAAT) platforms are currently available. Our laboratory uses two real time RT-PCR platforms, the Roche Cobas and the Cepheid GeneXpert Dx. Both platforms perform comparably in our experience, having similar limits of detection (LOD) of SARS-CoV-2 viral RNA, 100-200 and 250 copies/mL, respectively. However, the assay run times of 3.5 hours and 45 minutes, respectively, are still too long for timely decision support in a variety of important clinical situations (e.g, bed assignments for patients being admitted from the Emergency Department who may require cohorting by their COVID-19 status). In search for a platform with shorter turnaround time, we sought to evaluate the recently released Abbott ID NOW COVID-19 assay (ID NOW). The ID NOW assay uses isothermal nucleic acid amplification of the RdRp viral target with a claimed LOD of 125 genome equivalents /mL. The objective of our study was to evaluate the performance of the ID NOW test by using the Cepheid-Xpert-Xpress SARS-CoV-2 (Xpert Xpress) on the GeneXpert Dx as the comparator reference method.

## Materials and Methods

### Abbott ID NOW platform vs. Cepheid platform using nasopharyngeal swabs in VTM

All samples were obtained from patients who presented to the NYU Langone Hospitals (Manhattan) emergency department for whom a physician ordered a test for SARS-CoV-2. Nasopharyngeal (NP) samples were obtained using flocked NP swabs (NPS) and transported to the laboratory in universal viral transport media (VTM) (Cepheid Xpert Viral Transport Medium; SWAB/B-100) or BD Universal VTM / 2250527) within 1-2 hours of collection. All samples were NPS transported to the lab in VTM.

Initial verification of the ID NOW platform was accomplished by using two previously tested positive patients’ samples using the Xpert Xpress assay, one with a Ct value 30.8 for N2 and another with a Ct value of 19.3 (i.e. high viral load). Both samples were diluted 1:2, 1:5, 1:10, 1:20, 1:50 and 1:100 in VTM. All the dilutions were tested in replicates on both platforms.

We assessed the performance of Abbott ID NOW on 15 consecutive positive (by Xpert Xpress) NP specimens submitted in VTM from patients seen at the emergency department with suspected COVID-19 disease. Each specimen was tested on the ID NOW platform.

ID NOW performance in relation to N2 Ct values obtained by the Xpert Xpress assay was conducted by testing a set of 8 previously tested specimens using the Xpert Xpress assay that had increasing N2 Ct values ranging from 32.6 to 41.8. All samples were tested on the ID NOW and the results compared to those previously obtained using the Xpert Xpress.

In response to our initial findings of lower positive agreement of the ID NOW compared to the Xpert Xpress, Abbott communicated to us that despite the indications in the package insert, their assay was not optimized for testing NPS in VTM; the best performance would be from dry nasal swabs obtained and tested at the point of care (POC). They indicated that they would be modifying their package insert to reflect this change (ID NOW COVID-19 Product Insert, IN190000 Rev.3 2020/04:6-8).

### Abbott ID NOW platform using dry nasal swabs vs. Cepheid platform using nasopharyngeal swabs

We compared the Abbott ID NOW platform using paired dry nasal swabs (both nares) to the Cepheid platform using NPS in VTM. We collected paired nasal swabs and nasopharyngeal swabs in parallel from patients who presented to the NYU Langone Hospitals (Manhattan) emergency department for whom a physician ordered a test for SARS-CoV-2. Tests were ordered by the ED physician based on the NYU ED protocol for potential COVID-19. Collection instructions for the nasal swabs provided by Abbott were communicated to ED care providers. Nasal samples were obtained with swabs supplied with the Abbott assay (Puritan Medical Products 25-1506 IPF100). While the sequence of collecting both sample types varied, all NPS were obtained through one nostril and nasal samples were obtained from both nares. Nasopharyngeal swabs were collected as described above.

Although Abbott indicated that the dry swab could be transported in the same sleeve in which the swabs were packaged, we chose to transport the samples in a sterile tube in order to minimize loss of sample on the sleeve and ensure safety during transport. A total of 101 paired samples collected.

Dry nasal swabs were transported to the laboratory at room temperature in a sterile tube along with the corresponding NPS in VTM. The dry nasal swabs were tested within 2 hours of collection or kept refrigerated at 4-8°C for up to 24 hours before testing as per package insert instructions.

### Instruments

#### ID NOW COVID-19

(Abbott Diagnostics Scarborough, Inc., Scarborough, ME), is a rapid test that qualitatively detects SARS-CoV-2 viral nucleic acids from nasal, nasopharyngeal and throat swabs. It is an automated assay that utilizes isothermal nucleic acid amplification technology. The assay amplifies a unique region of the RdRp genome with a manufacturer’s claimed LOD of 125 genome equivalents/mL. Positive results are available within 5-13 minutes and negative results within 13 minutes.

#### Xpert Xpress SARS-CoV-2 test

(Cepheid, Sunnyvale, CA) is a rapid, real-time RT-PCR test that detects SARS-CoV-2 RNA in nasopharyngeal swab and/or nasal wash/aspirate specimens. The assay amplifies 2 nucleic acid targets, namely N2 (nucleocapsid) and E (envelope) wherein N2 is more specific for SARS-CoV-2. The LOD for this assay is claimed by the manufacturer to be 250 copies/mL. SARS-CoV-2 genomic load is assessed using the number of amplification cycles needed for a positive PCR test (i.e. the cycle threshold [Ct] value). Ct values have an inverse relationship to viral loads and thereby provide a surrogate measurement of the viral genomic load (7). The Cepheid GeneXpert instrument interprets the results automatically based on detection of N2 and E targets. For a positive detection, the Ct values must be within a valid range and end-point above the minimum setting as determined by the manufacturer software. Both assays are used only under Food and Drug Administration’s Emergency Use Authorization (EUA) (8).

The performance evaluation of the Abbott ID NOW assay was done as a routine laboratory quality assessment as normally conducted before introducing any new assay.

### Statistical analysis

Qualitative Method Comparison was performed using EP Evaluator (Data Innovations, VT 05403), Evidence-Based Medicine Diagnostic Toolbox (**https://ebm-tools.knowledgetranslation.net/calculator/diagnostic/)** and the correlation coefficient using Excel 2010.

## Results

### Abbott ID NOW platform vs. Cepheid platform using nasopharyngeal swabs in VTM

Results of the initial verification of the ID NOW revealed that there was good positive percent agreement (PPA) with Xpert Xpress when a specimen (Sample 1) with low N2 CT value (19.3) was progressively diluted up to 1:100. However, this was not the case for a specimen (Sample 2) with initial higher N2 Ct value (30.8) (Table 1). Lower positive percent agreement between the two platforms was noted at 1:10 dilution level and beyond with only a 33% agreement at 1:100 dilution.

**Table 1:** Initial verification of ID NOW. Positive Agreement* when testing two progressively diluted patient specimens with low and high Xpert Xpress N2 Ct values

| Patient Specimen** | Dilution | Replicate runs | Abbott Results in Agreement with Reference Method |
| --- | --- | --- | --- |
| Sample 1 | 1:2 | 1 | 1 (100%) |
| Sample 1 | 1:5 | 1 | 1 (100%) |
| Sample 1 | 1:10 | 4 | 4 (100%) |
| Sample 1 | 1:20 | 3 | 3 (100%) |
| Sample 1 | 1:50 | 2 | 2 (100%) |
| Sample 1 | 1:100 | 2 | 2 (100%) |
| Sample 2 | 1:2 | 1 | 1 (100%) |
| Sample 2 | 1:5 | 1 | 1 (100%) |
| Sample 2 | 1:10 | 4 | 3 (75%) |
| Sample 2 | 1:20 | 4 | 2 (50%) |
| Sample 2 | 1:50 | 3 | 2 (67%) |
| Sample 2 | 1:100 | 3 | 1 (33%) |

\* All samples dilutions were detected by Xpert Xpress
\*\*Sample 1 Xpert Xpress N2 Ct: 19.3 (undiluted). Sample 2 Xpert Xpress N2 Ct 30.8
(undiluted).

Comparison of results from our first study of Abbott ID NOW obtained on 15 consecutive NP positive specimens in VTM that was previously tested by Xpert Xpress produced 5 negative results by ID NOW (PPA of 66.7%). The Xpert Xpress N2 Ct values of the discordant specimens ranged from 36.3 to 44.3.

In order to confirm that the lower sensitivity of the ID NOW platform was related to the Ct value obtained on the Xpert Xpress, we tested a series of previously positive NP specimens with increasing N2 Ct values. Similar to the dilution studies, the performance of the ID NOW started to decline as the Xpert Xpress N2 Ct values increased throughout the series, although at a higher Ct value (38.0) than that seen with the earlier tested consecutive positive specimens from ED patients (36.3).

The distribution of the Xpert Xpress Ct values of 1439 positive samples submitted from different areas of our institution in the previous six weeks revealed that 15.1% of total positive samples had Ct values above 38 (data not shown), the percentage increasing over more recent weeks.

### Abbott ID NOW platform using dry nasal swabs vs. Cepheid platform using nasopharyngeal swabs

For the comparison of dry nasal swabs versus NP samples in VTM, with testing in ID NOW and Xpert Xpress, respectively, we included samples from 101 ED patients (age 28 - 90) with all suspected of COVID (onset of symptoms 1 day to 1 month) except one patient who was tested prior to surgery for ectopic pregnancy as per emergency department protocols. As seen in Table 2, the ID NOW using dry nasal swabs agreed in 17 of 31 positive Xpert Xpress samples, PPA of 54.8% (95% CI: 37.8-70.8%). The remaining 14 samples were not detected by ID NOW. The ID NOW results matched 69 of the 70 negative Xpert Xpress results with one detected by ID NOW but not by Xpert Xpress, Negative Percent Agreement (NPA) 98.6% (95% CI:92.3 - 99.7%). Overall, there was 85.1% Agreement (95% CI 76.9 - 90.8%), and a Positive Predictive Value (PPV) and Negative Predictive Value (NPV) of 94.4% (95% CI:74.3 - 99.0 %) and 83.1% (95% CI :73.7 −89.7%) respectively.

**Table 2.**
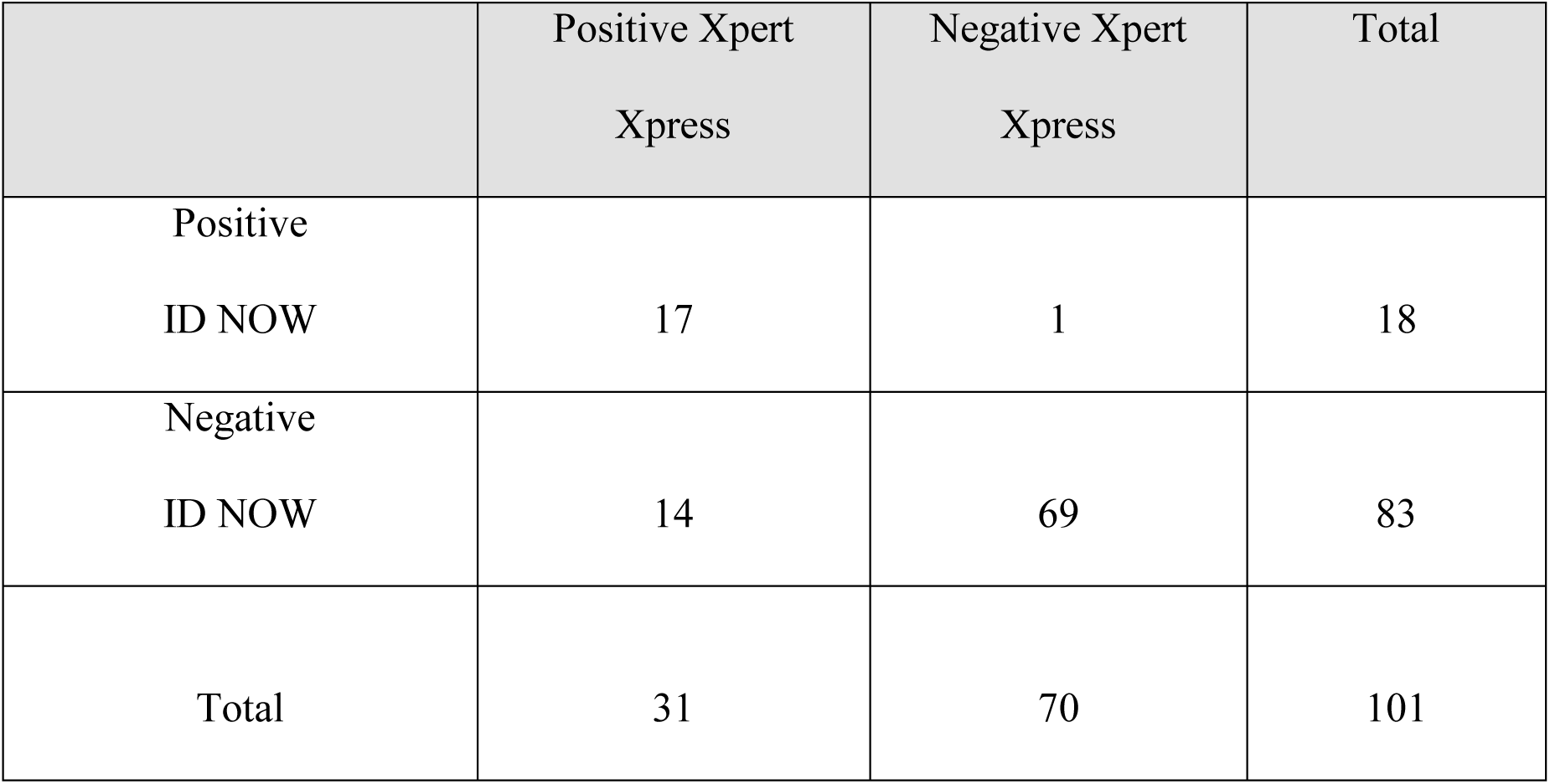
Comparison of ID NOW using dry nasal swabs and Xpert Xpress using NPS in VTM on 101 paired samples.

|  | Positive Xpert<br>Xpress | Negative Xpert<br>Xpress | Total |
| --- | --- | --- | --- |
| Positive<br>ID NOW | 17 | 1 | 18 |
| Negative<br>ID NOW | 14 | 69 | 83 |
| Total | 31 | 70 | 101 |

Grouping the Xpert Xpress N2 Ct values arbitrarily into low, medium and high ranges showed that all 6 dry nasal swabs with their corresponding NP in VTM having a N2 Ct value of <33.5 by Xpert Xpress tested positive by ID NOW. But the PPA of ID NOW was 54.5% (6 of 11) and 35.7% (5 of 14) when N2 Ct values obtained by Xpert Xpress were between 33.7 and 38, and >38, respectively.

The average time between collection and testing of the samples with discrepant results and those testing positive by the ID NOW was approximately 90 minutes.

There was no correlation between N2 Ct and days of onset of symptoms (*R2* = 0.097). Nineteen of the 31 patients whose samples tested positive for SARS-CoV2 RNA by Xpert Xpress were admitted. Of the 14 “ID NOW negative/ Cepheid Xpert Xpress positive” patients one was admitted to hospice and 5 were admitted to NYU Langone Health - Tisch Hospital; one of these patients was admitted for ectopic pregnancy surgery. Thirteen of 14 patients whose samples were ID NOW negative/ Cepheid Xpert Xpress positive had clinical or radiological evidence of COVID.

## Discussion

The urgency of the COVID-19 pandemic and the United States government’s subsequent use of the EUA has placed an unusual onus of responsibility on the clinical laboratories to substantiate the clinical value and performance of the COVID-related tests they introduce. Accurate diagnostic testing and subsequent isolation of infected individuals is crucial to mitigating continued transmission. The appeal of the Abbott technology stemmed from our intention to provide rapid and accurate results in our institution, especially for our Emergency Department.

Our results showed progressively lower PPA with lower viral load (higher N2 Ct values) as determined by Xpert Xpress, but this was not consistent. In our comparison studies of paired dry nasal swabs and NP swabs in VTM, there were samples testing positive by Abbott platform even when Xpert Xpress N2 Ct values were above 38 which might suggest better sampling using the dry swab in these cases. A study by Zhen et al. has also shown discordant results comparing these two assays against the Hologic Panther Fusion SARS-CoV-2 assay as the reference method (9). The ID NOW produced 7 false negatives out of 57 positive NP samples, resulting in a PPA of 87.7% compared to 98.3% on Xpert Xpress. Their study however, included a total of 108 NPS transported in VTM, which the manufacturer has now noted decreases the sensitivity of their assay. These authors also determined that the Xpert Xpress had the lowest LOD at 100 viral copies/mL, while the ID NOW had an LOD of 20,000 copies/mL (9), well above the limit of detection stated in the Abbott ID NOW COVID-19 package insert (125 genome equivalents /mL).

Smithgall et al., found an overall PPA of 73.9% with ID NOW, 98.9% with the Xpert Xpress when compared against the Cobas Roche assay (10). Similar to our study, these authors found that the performance of the ID NOW decreased for samples with high Ct values. The study by Rhoads on 96 NPS, however, found a PPA of 94% for ID NOW, compared to 96% with Simplexa Diasorin against a modified CDC assay. This study also showed a correlation with Ct values (R2=0.89 - 0.91). This would suggest that the sensitivity of both the ID NOW and the Simplexa was reduced as the Ct values exceeded 34.5 (11).

Although the original Abbott package insert had stated that 0.5 to 3.0 mL of viral transport media was acceptable for use in their assay, it also recommended minimizing dilution of the sample as it might result in decreased test sensitivity. The use of dry swabs was recommended by the manufacturer for optimal test performance. Contrary to manufacturer expectations, however, our parallel study showed that the PPA of ID NOW using dry nasal swabs (54.8%), was actually lower than when using NPS in VTM (66.7%) on 15 consecutive positive samples as determined by Xpert Xpress in this study.

Interestingly, a recent study by Harrington et al. compared the results of 524 dry nasal swabs tested on the ID NOW with paired nasopharyngeal swabs in VTM run on the Abbott m2000 RT-PCR system (12). The specimens were submitted from 3 Emergency Departments and 2 Immediate Care Centers; about two thirds of the dry swabs were tested at the point of care. The ID NOW showed an overall 74.7% positive agreement with the m2000 and 70.6% for the group tested at the point of care.

The literature provides no strong consensus regarding the influence of transport medium and storage conditions on viral load stability. One study looking at differing viral transport media and storage conditions using the Quest and Roche EUA assays showed consistent detection of SARS-CoV-2 RNA after 7 days at room temperature, and up to 14 days refrigerated or frozen (13). The ID NOW package insert also directs clinicians to test swabs as soon as possible after collection, and if not possible, to hold it in its original package for no more than two hours prior to testing or to refrigerate up to 24 hours prior to testing. Outside of inpatient hospital laboratories, such as in outpatient point-of-care testing, independent validation should be performed to assess performance in different settings, including pre-analytic variables.

Extrapolating from our results from 101 patient samples, if the ID-NOW assay were to be used as a first step screening, confirmation of over 80% of the tested samples would be required to be confident that the negative results are truly negative. While there was one apparent false-positive result in our parallel study, the overall high PPV (94.4%) and NPA (98.6 %) do support the added value that in 5 −13 minutes a positive result can be interpreted as a true positive. We could not confirm that the apparent false positive result obtained with the Abbott ID NOW was due to sampling variability.

Based on our findings, the ID NOW has some utility as a rapid rule-in test for COVID-19 with samples at high viral load; however, we advise caution with its use as a singular rule-out test especially in the setting of samples with lower viral loads. Of the positive samples missed by ID NOW, 13 of the 14 patients had clinical or radiologic evidence of COVID-19. The remaining patient was an asymptomatic patient admitted for an ectopic pregnancy. Currently we cannot assume that patients with low viral load are unable to transmit the virus, as a false negative can significantly affect subsequent clinical and epidemiologic decision-making. It is noteworthy that ID NOW exhibited higher target detection when using NPS in VTM as compared with the dry nasal swabs. Patients having lower levels of SARS-CoV-2 virus in their nasopharynx might harbor even lower concentrations in their nasal cavity leading to a lower level of detection by the ID NOW.

Reasons for the lower PPA between the ID NOW and Xpert Xpress could be attributed to several factors. The sample population included in our study during the latter part of April 2020, although having a broad spectrum of Xpert Xpress-determined N2 Ct values, did have a significant number of samples with Ct values above 38, which ID NOW did not detect consistently. Differences in viral RNA detection could be related to the differences in NAAT technology used. Although isothermal nucleic acid amplification test (INAAT) is a promising tool, it has been found to have lower sensitivity compared with RT-PCR for SARS-CoV-2 RNA detection (14). Compared to RT-PCR, the molecular chemistry of INAAT is more complex, involving more primers and intricate conformational rearrangements through enzymatic strand displacement, emphasizing the importance of target selection and primer design on performance.

Another possible explanation is the potentially poorer performance of the RdRp target used in the Abbott assay compared to other RNA targets as has been described (15, 16). Recent reports indicate that mutations in the RdRp domain have been emerging (17). It is unknown at this point if the genomic region targeted by the Abbott assay has changed significantly, as the virus has spread throughout the world.

Limitations of our study include relatively small sample size, inability to control for sampling variability and lack of an additional comparator method to discern the discrepancies. In our laboratory, we periodically perform correlation studies between our 2 platforms, Cobas and Cepheid, and results consistently match. Another limitation was transporting the dry swabs specimens to the laboratory for testing rather than testing directly in a point-of-care setting as the Abbott ID NOW platform is intended. Nevertheless, this factor might not have influenced our results, since the samples were handled and tested in ID NOW within the appropriate time and conditions stipulated in the package insert.

Overall, our study revealed low PPA of ID NOW when compared with Xpert Xpress irrespective of use of viral transport media or sample type, which raises concerns regarding its suitability as a diagnostic tool. An assay that detects a broad range of the SARS-CoV-2 RNA levels is highly desirable since it is difficult to control the sample quality and as a consequence, variations in viral concentrations. At this point, the significance of low viral levels in disease or transmission is largely unknown.

## Acknowledgements

We sincerely acknowledge the help and support of the medical technologists from the Tisch Microbiology laboratory (Jiamin Chui, Claudia Funez and Alicia Lovell) and the Emergency Medicine Nursing Team including Patricia Baptiste, Jennifer Tiarsmith, Jason Garcia, Roselle David, Robin Keida, Jacquelyn Burns, Lauryn Clouden for caring for the patients and their assistance in collecting the samples used in this study.

